# Measuring *C. elegans* spatial foraging and food intake using bioluminescent bacteria

**DOI:** 10.1101/759928

**Authors:** Siyu Serena Ding, Karen S. Sarkisyan, Andre E. X. Brown

## Abstract

For most animals, feeding includes two behaviours: foraging to find a food patch and food intake once a patch is found. The nematode *Caenorhabditis elegans* is a useful model for studying the genetics of both behaviours. However, most methods of measuring feeding in worms quantify either foraging behaviour or food intake but not both. Imaging the depletion of fluorescently labelled bacteria provides information on both the distribution and amount of consumption, but even after patch exhaustion a prominent background signal remains, which complicates quantification. Here, we used a bioluminescent *Escherichia coli* strain to quantify *C. elegans* feeding. With light emission tightly coupled to active metabolism, only living bacteria are capable of bioluminescence so the signal is lost upon ingestion. We quantified the loss of bioluminescence using N2 reference worms and *eat-2* mutants, and found a nearly 100-fold increase in signal-to-background ratio and lower background compared to loss of fluorescence. We also quantified feeding using aggregating *npr-1* mutant worms. We found that groups of *npr-1* mutants first clear bacteria from each other before foraging collectively for more food; similarly, during high density swarming, only worms at the migrating front are in contact with bacteria. These results demonstrate the usefulness of bioluminescent bacteria for quantifying feeding and suggest a hygiene hypothesis for the function of *C. elegans* aggregation and swarming.

## INTRODUCTION

Feeding behaviour plays an important role in fields ranging from ecology and evolution (Larsen 2003; MacArthur and Pianka 1966) to ageing and metabolism (Balasubramanian, Howell, and Anderson 2017; Trepanowski et al. 2011) and health and disease (Djalalinia et al. 2015; Mattson et al. 2014). The roundworm *C. elegans* has emerged as a useful model organism to study all aspects of feeding, including worms’ immediate response to finding food (Sawin, Ranganathan, and Horvitz 2000), foraging and patch leaving (Shtonda 2006; Harvey 2009; Bendesky et al. 2011; Milward et al. 2011; E. Scott et al. 2017), as well as the details of food intake (L. Avery 1993; Leon Avery and Shtonda 2003; Fang-Yen, Avery, and Samuel 2009) and even spitting (Bhatla et al. 2015).

These studies of the genes and neural circuits underlying feeding rely on a variety of methods that have been developed to quantify feeding in *C. elegans. C. elegans* feeds by sucking bacteria into its mouth using rhythmic pumping of pharynx (Avery and You 2012), and pharyngeal pumping frequency is often used as a proxy for food intake. Because worms are transparent, pharyngeal pumping can be measured manually by direct observation under a stereomicroscope or more recently, using automated image analysis (Scholz et al. 2016). Electrophysiological readouts can also be used to measure multiple worms in parallel in microfluidic devices (Lockery et al. 2012). Alternatively, feeding can be measured using a non-food additive such as exogeneous luciferin (Rodríguez-Palero et al. 2018), dye (You et al. 2008), or fluorescent beads (Fang-Yen et al. 2009; Kiyama et al. 2012). Bacteria consumption can also be measured directly by optical density in liquid (Gomez-Amaro et al. 2015) or by using fluorescently-labelled bacteria. Labelled bacteria can provide a quantitative measurement of food inside the worm gut using a worm sorter (Andersen et al. 2014) or image analysis (You et al. 2008), and consumption can be measured on solid media using a plate reader (Zhao et al. 2018).

How food is distributed and consumed in space has crucial implications for animal foraging strategy (Bernstein 1975; Ding, Muhle, et al. 2019; Lanan 2014; Stenberg and Persson 2005) and subsequent fitness. Therefore, of the existing methods of quantifying feeding, imaging the consumption of fluorescently-labelled bacteria is of particular interest since it can provide information on both where and how much food has been consumed. However, as fluorescent proteins form stable cooperatively folding structures, they are resistant to proteolytic cleavage (Nicholls and Hardy 2013; Bokman and Ward 1981). This results in high background fluorescence signal even after bacteria are digested by *C. elegans*, complicating both the quantification and the interpretation of feeding behaviour.

Here we use an *E. coli* strain with self-sustained bioluminescence to monitor both the rate of food intake and its spatial distribution in laboratory reference and mutant worms, worms treated with serotonin and naloxone, and in high-density worm swarms.

## RESULTS

### Bioluminescent bacteria improve signal to background ratio in a feeding assay

Previous studies have used fluorescent protein-expressing strains of *E. coli* to measure worm feeding. However, when we recorded worms feeding on *E. coli* strain OP50-DsRed, we noticed a prominent background fluorescence signal, which was especially conspicuous in our experiments with DA609 (*npr-1* aggregation mutant) worms (Figure 1A, red arrow). These worms first form aggregates on food and then collectively swarm over the food patch following local food depletion (Ding, Schumacher, et al. 2019). It is possible that a fraction of DsRed molecules survives passage through the worm gut due to resistance to protease cleavage. Background signal may also result from fluorescent protein molecules that seeped into the medium from the cytoplasm of dead bacterial cells or have been expelled with liquid as a normal part of pharyngeal pumping. Alternatively, the background may be attributable to a small number of bacteria at a density that is low enough for worms to ignore, although this seems unlikely given that the background “halo” can be quite bright (Figure 1A, red arrow). These three possible sources of fluorescence background are not mutually exclusive and complicate the quantification and interpretation of feeding experiments. The background remained when we used a different fluorophore (*E. coli* OP50-GFP, Figure 1B, middle) or a different worm strain that does not aggregate (*C. elegans* N2, Figure 1B, right).

**Figure 1.**
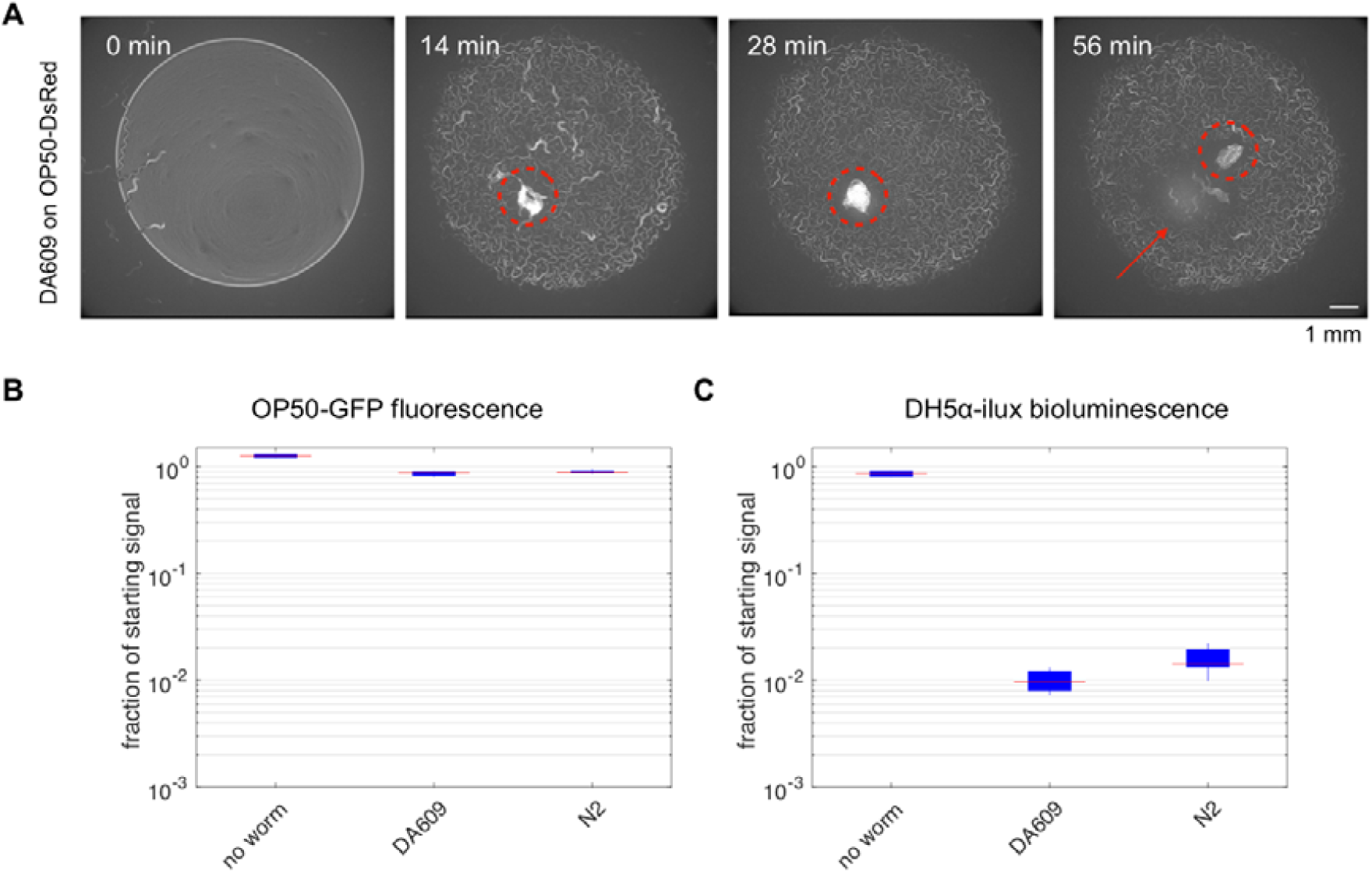
Assessing worm feeding behaviour with bacteria labelling. **A**) Sample snapshots of a group of 40 DA609 worms feeding on a fluorescent *E. coli* OP50-DsRed lawn. Red circles show the worm aggregate, and the red arrow points to the remaining background signal after the worm cluster moves away from its original site. **B-C**) Background signal comparison between fluorescence (B) and bioluminescence (C) methods. Population feeding experiments were performed on low peptone NGM plates seeded with fluorescent *E. coli* OP50-GFP (B) or bioluminescent *E. coli* DH5α-ilux (C). Zero (“no worm”) or 40 (“DA609” and “N2”) worms were allowed to feed on and deplete the bacteria for 13.5 hours. Signal from the labelled bacteria was obtained at the start (100% food) and at the end (0% food) of the experiment using the fluorescence (465 nm excitation, 520 nm emission) or the bioluminescence (no excitation, open emission) imaging protocol, and the final to starting signal ratios were calculated. n = 2 for no-worm, n = 6 for DA609, n = 6 for N2, pooled between two independent sets of experiments.

As an alternative to fluorescence-based bacterial labelling, we tested a bioluminescent *E. coli* strain (DH5α-ilux) as the worm food source. We transformed *E. coli* DH5α with a plasmid containing an engineered *Photorhabdus luminescens lux* operon encoding enzymes of a bacterial bioluminescence system (Gregor et al. 2018). These enzymes perform biosynthesis, oxidation and recycling of a long-chain fatty aldehyde, the key component of the light-emitting reaction along with flavin mononucleotide. We seed a defined quantity of DH5α-ilux liquid culture onto Nematode Growth Medium (NGM) plates, let a population of 40 worms feed, and monitor food consumption over time using an IVIS Spectrum imaging system. We show that following DA609 or N2 feeding experiments that result in total food exhaustion, the bioluminescence imaging method gives very reduced background when normalised against the starting signal (Figure 1C), in contrast to fluorescence imaging which shows noticeable background levels (Figure 1B). Feeding assays using bioluminescent bacteria shows a nearly 100-fold increase in signal-to-background ratio compared to using fluorescent bacteria (Figure 1B-C).

### Bioluminescence depends on growth conditions and provides a quantitative measurement of worm feeding rates

Since bioluminescence from DH5α-ilux depends on active bacterial metabolism, we next characterised signal strength under different experimental conditions. Storing bacterial culture at 4°C overnight abolishes the signal, so a fresh overnight culture was prepared for all experiments. We grew DH5α-ilux in overnight liquid cultures at 37°C to stationary phase and allowed them to cool down to room temperature before inoculating onto NGM media for imaging. Serial dilution of the overnight liquid culture shows roughly linear scaling with bacteria concentration (Figure 2A). After inoculating 20 μL of overnight culture onto NGM media plates containing different levels of peptone (regular peptone, 0.25% w/v; low peptone, 0.013% w/v; no peptone, 0% w/v), bioluminescence signal was monitored for hours (Figure 2B) and days (Figure 2C) at 20°C. As expected, signal is the highest on media with the highest peptone concentration (Figure 2B-C, blue lines) and lowest on no-peptone media (Figure 2B-C, black lines). On standard NGM (0.25% peptone), bioluminescence increases for approximately one week and then decreases over several days with no obvious stationary plateau (Figure 2C, blue line); On the scale of hours, there is an initial decrease in intensity over the first few hours followed by an approximately 5-fold increase over the next day (Figure 2B, blue line). This initial decrease perhaps represents a lag phase of growth on solid media. Therefore, bioluminescence signal strength from DH5α-ilux depends on a number of growth conditions that affect bacterial metabolism, including temperature, peptone level, and inoculation time.

**Figure 2.**
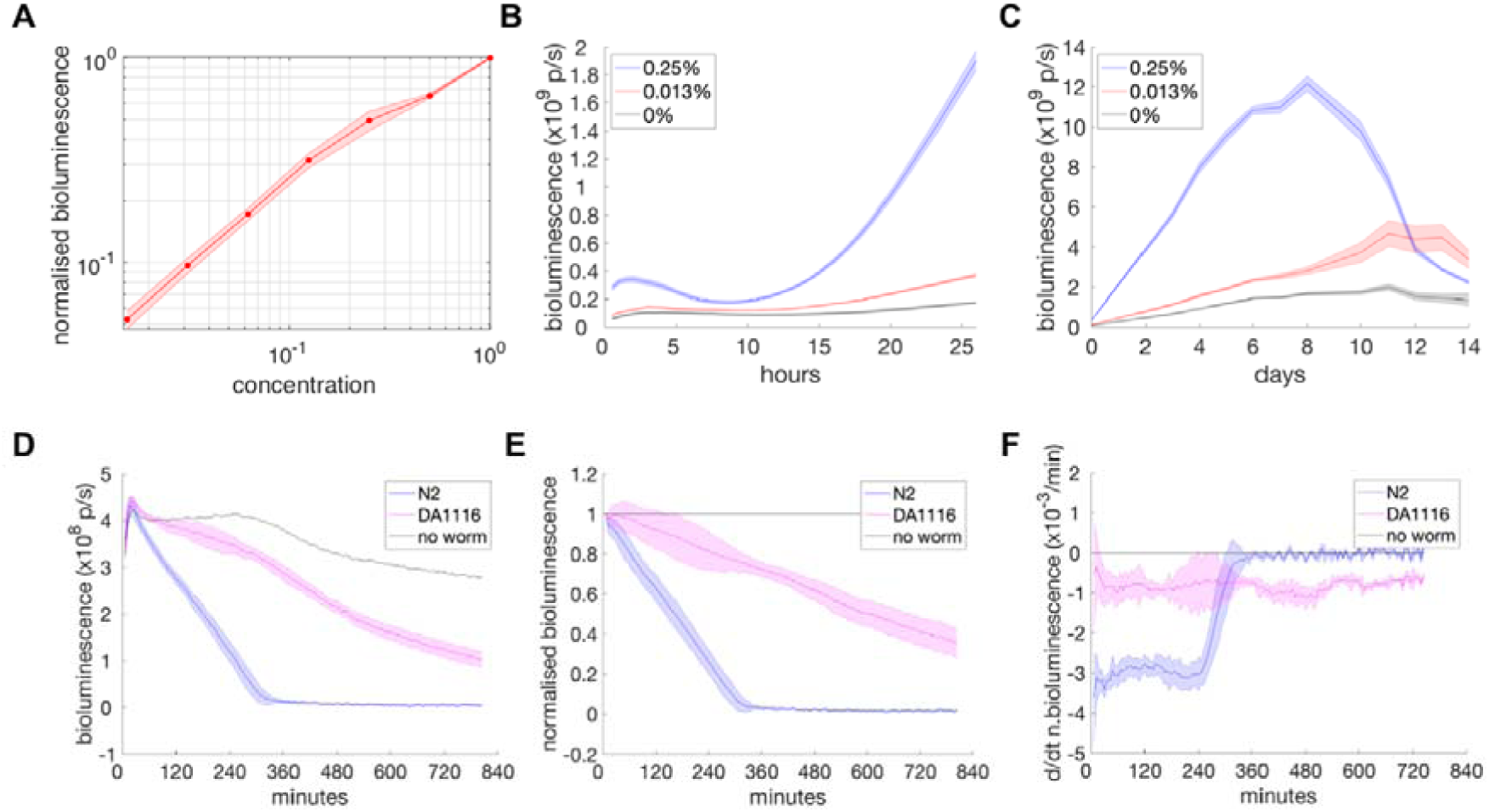
DH5α-ilux bioluminescence signal characterisation. **A**) Normalised signal from liquid bacteria culture in a 2-fold dilution series. Concentration of 1 is undiluted bacteria overnight culture. Signal is taken immediately following serial dilution in LB broth where all samples have a final volume of 150 μL, and are normalised to undiluted levels. Here n = 6, pooled between two independent sets of experiments. Error bars represent ±1 standard deviation (SD). **B-C**) Signal from 20 μL of bacteria culture after hours (B) and days (C) of inoculation on NGM media containing different peptone levels. “p/s” is photons/s. All inoculations were performed at 20 °C, and all measurements were made following a 1 second exposure. For B), signal was taken every 30 minutes and all samples were imaged simultaneously. n = 3 for each condition, error bars represent ±1 SD. For C), signal was taken once on most days, n = 3, error bars represent ±1 SD. **D-F**) Bioluminescence from population feeding experiments of N2 and DA1116 worms, showing **D**) raw signal, **E**) normalised signal (normalised against the starting signal and then against the control signal), **F**) derivative of the normalised signal calculated over a 60-minute window). Forty N2 (blue) worms, forty DA1116 (magenta) worms, or no-worm control (black) experiments were performed on a 20 μL DH5α-ilux lawn. Measurements were taken every 6 minutes using 1 second exposure. All samples shown were imaged simultaneously. Here n = 6 for N2, n = 4 for DA1116, n = 2 for control, pooled from two independent sets of experiments; error bars represent ±1 SD.

We next compared the population feeding rates of the laboratory reference N2 strain and DA1116, an *eat-2* mutant with abnormal neurotransmission in the pharynx (McKay et al. 2004) and that pumps slowly (Raizen, Lee, and Avery 1995). To take into account different initial bioluminescence levels across experimental samples (Figure 2D), we divide the signal in each condition by the level detected in the first frame. This relative signal is then further normalised by the value of the corresponding no-worm control at each time point to correct for the non-stationarity of the signal in the absence of feeding (Figure 2E). Relative feeding rates are then estimated by taking the derivative of the normalised signals over time (Figure 2F). Since the normalisation is important for reliably estimating the feeding rate, we recommend including no-worm controls whenever possible.

We show that both N2 and DA1116 worm strains deplete the food at a roughly constant rate (Figure 2E-F) and that the median feeding rate from the first four hours (before N2 runs out of food) for DA1116 is 27% that of N2. DA1116’s reduced feeding rate on solid media is consistent with previous reports of its slow pumping (∼10% that of N2 (Raizen, Lee, and Avery 1995)) and restricted food intake in liquid-based assays (∼80% that of N2 as measured by optical density-based bacterial clearing (Gomez-Amaro et al. 2015) and ∼60% that of N2 as measured by luciferin ingestion (Rodríguez-Palero et al. 2018)). This experiment also confirms that the signal from freshly inoculated overnight liquid culture is sufficient to estimate relative feeding rates. For less sensitive imaging instruments, it would be possible to incubate seeded plates for longer to obtain higher signal (Figure 2C).

We repeated the feeding experiments and analysis using OP50-GFP bacteria (Supplementary Figure S1) instead of DH5α-ilux, and obtained similar results showing that DA1116 feeding rate is 27% that of N2 despite a very different no-worm control signal (Supplementary Figure S1A, black line). This highlights the importance of normalisation using either bacteria labelling method. While the relative feeding rate results are reassuringly similar between bioluminescence- and fluorescence-based methods, the fluorescence method shows a lower signal-to-background ratio as well as high background levels (Supplementary Figure S1A) that may complicate analysis and interpretation when normalised (Supplementary Figure S1B-C).

Serotonin has previously been reported to enhance pharyngeal pumping and food intake (Horvitz et al. 1982; Niacaris and Avery 2003) as well as the slowing response of starved worms (Sawin, Ranganathan, and Horvitz 2000). We thus pre-starved N2 worms before exposing them to serotonin in the presence of food. Unexpectedly, serotonin-treatment caused a decrease in the measured feeding rate (Supplementary Figure S2A, blue and black lines). We observed a comparable reduction in feeding rate using OP50-GFP bacteria as the food source (Supplementary Figure S2B, blue and black lines). We confirmed serotonin was having the expected effects on pumping rate (Supplementary Figure S2D). However, in serotonin-treated samples, a smaller number of worms reaches the bacterial lawn (Supplementary Figure S2E-F), most likely due to serotonin’s suppression of locomotion (Horvitz et al. 1982). Therefore, these results do not contradict previous findings but highlight the multifaceted effects of serotonin and the essential role of foraging in successful feeding.

In contrast to serotonin, the morphine antagonist naloxone has been reported to decrease food intake in starved worms by acting on an opioid receptor expressed in a sensory neuron (Cheong et al. 2015). Naloxone treatment did result in a decreased feeding rate as expected (Supplementary Figure S2A, red and black lines), but the levels of bioluminescence were much lower than in naloxone-free controls (Supplementary Figure S2C, left). We again confirmed that the feeding rate was reduced using OP50-GFP bacteria (Supplementary Figure S2B, red and black lines). Fluorescence levels were decreased compared to controls (Supplementary Figure S2C, right) but the effect was not drastic and naloxone does not detectably affect *E. coli* growth (Maier et al. 2018), suggesting that naloxone may act more specifically on bioluminescence-related metabolism.

### Bioluminescent bacteria reveal the spatial aspect of *C. elegans* feeding behaviour

To gain insights into different *C. elegans* feeding strategies, we performed experiments with the laboratory reference strain N2 and DA609, an *npr-1* loss-of-function mutant. The former are solitary feeders whereas the latter are social, initially forming worm aggregates on food and then collectively swarming over the food patch following local food depletion (Ding, Schumacher, et al. 2019). Our feeding experiments on DH5α-ilux bioluminescent bacteria show that DA609 and N2 populations both have stable feeding rates, and that DA609 has a feeding rate that is 58% that of N2 (Figure 3A-B). Experiments with OP50-GFP fluorescent bacteria also show a relative feeding rate of 58% (Supplementary Figure S3). Thus the social feeders have a lower feeding rate than the solitary ones despite similar pharyngeal pumping rates (Choi et al. 2013), consistent with a previous report measuring the amount of fluorescently labelled bacteria inside worm guts (Andersen et al. 2014).

**Figure 3.**
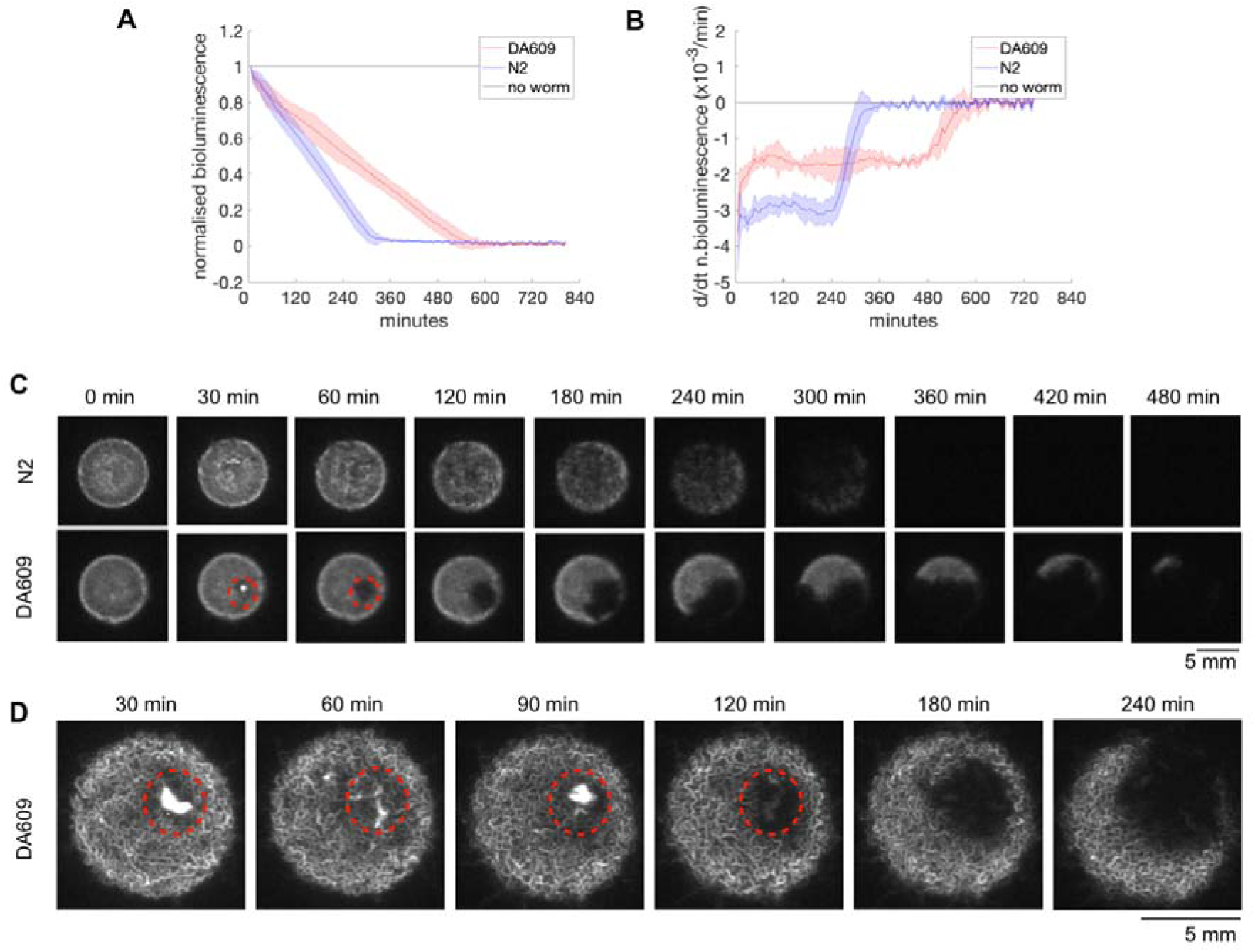
Bioluminescence from population feeding experiments of N2 and DA609 worms, showing **A**) normalised signal, and **B**) derivative of the normalised signal calculated over a 60-minute window. Forty DA609 (red) or N2 (blue) worms or no-worm control (black) experiments were performed on a 20 μL DH5α-ilux lawn. One second exposure measurements were read every 6 minutes. n = 6 for each condition, pooled between two independent sets of experiments; error bars represent ±1 SD. **C**) A series of snapshots contrasting the spatial pattern of food depletion in N2 (top) and DA609 (bottom) population feeding experiments. **D**) A series of snapshots showing a DA609 worm aggregate (red circles) depleting food within the cluster first before moving onto new food.

Bioluminescence imaging also provides spatial information and we examined the pattern of food depletion between the two feeding strategies and noted major differences. While N2 worms show gradual depletion of the whole food patch roughly uniformly (Figure 3C, top row; Supplementary Movie S1, middle row), DA609 worms deplete food in a highly localised manner starting at one point and sweeping over the surface (Figure 3C, bottom row; Supplementary Movie S1, top row). These foraging behaviours observed here by bacterial depletion are consistent with our previous results in which worms were imaged directly (Ding, Schumacher, et al. 2019).

Moreover, we noticed that when DA609 worms initially aggregate they are covered in bacteria (Figure 3C-D, 30 min panels) and that the cluster stays in roughly the same place (Figure 3C-D, red circles) until the in-cluster bacteria are completely consumed. This observation fits well with the distinct “aggregation” versus “swarming” phases that we previously reported for DA609 (*npr-1*) aggregation (Ding, Schumacher, et al. 2019), suggesting that minimal cluster movement during the “aggregation” phase is due to the initial food availability inside the cluster. By contrast, the total depletion of bacteria inside the aggregate before collective movement starts is difficult to detect from the recordings of worms feeding on fluorescent bacteria, because the moving worm cluster is still fluorescent (Figure 1A, last panel; note the aggregation timescale is different for this experiment because OP50-DsRed bacteria were diluted). As mentioned previously, the source of this signal is unknown, but the bioluminescence results suggest that it is not due to residual metabolically active bacteria that adhere to the worm surface.

### Large-scale *C. elegans* swarms form a stable moving front on food

To study the behaviour of larger populations of swarming worms, we imaged thousands of young adult N2 worms feeding on larger 500 μL patches of DH5α-ilux bacteria and observed coherent swarming as the migrating worm front consumes the bacterial lawn in a single pass (Figure 4A; Supplementary Movie S2). Similar results were seen using OP50-GFP, although the fluorescent background remains after the front has passed (Supplementary Figure S4A; Supplementary Movie S3). Bacterial signal during swarming can be quantified using the same analysis methods as in 40-worm feeding experiments (Figure 4B-C).

**Figure 4.**
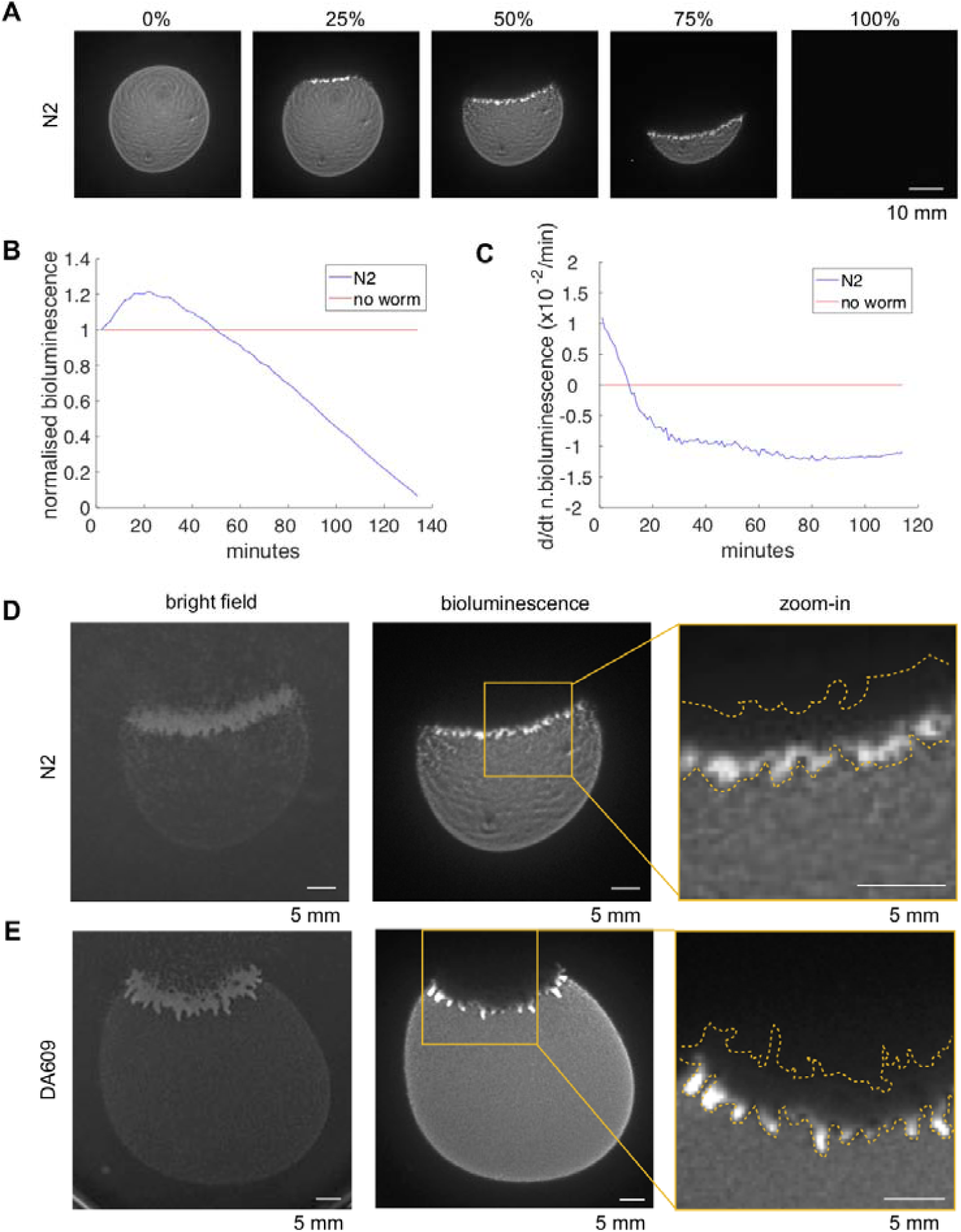
Bioluminescence signal from large population swarming experiments. A few thousand age-synchronised worms were allowed to feed and swarm over a 500 μL DH5α-ilux lawn. **A**) Snapshots of N2 swarming experiments, with time progression to total food depletion indicated at the top. **B)** Normalised signal and **C)** derivative of the normalised signal calculated over a 20-minute window, from an N2 swarming experiment with 1 day-old DH5α-ilux lawn. **D-E**) Sample snapshots from N2 (D) and DA609 (E) swarming experiments, showing bright field (left) and bioluminescence (middle) channels. The boxes in the middle panels are zoomed in and displayed on the right, with the worm front outline shown in dashed yellow lines.

Large populations of DA609 worms also swarm (Figure 4E). By overlaying the bioluminescence channel (Figure 4D-E, middle) with the bright field images (Figure 4D-E, left), it is clear that only worms at the leading edge of the migrating front are in contact with bacteria regardless of the worm strain (Figure 4D-E, right). We confirmed these results using OP50-GFP bacteria (Supplementary Figure S4), although the bacterial gradient is less obvious due to background fluorescence. Our results are similar to those in a recent study reporting a bacterial gradient in swarming *C. elegans* using OP50-GFP (Demir, Yaman, and Kocabas 2019). Finally, DA609 swarms form pronounced finger-like projections at the leading edge of the migrating front that protrude into the bacterial lawn (Figure 4E; Supplementary Movie S4; Supplementary Figure S4E).

## DISCUSSION

We have developed *C. elegans* feeding assays using bioluminescently labelled bacteria. This method allows simultaneous quantification of food intake and visualisation of food distribution, which are both important aspects of *C. elegans* feeding behaviour even though food intake has previously received greater attention. We show that a bioluminescence-based method results in higher signal-to-background ratios that simplify analysis and interpretation compared to fluorescence-based methods. In addition, it circumvents issues associated with fluorescence imaging such as phototoxicity, bleaching, autofluorescence, and behavioural modulation. Compared with other imaging-based methods, our method directly measures the ingestion of bacteria, rather than estimating bacterial uptake using exogeneous dye (You et al. 2008), beads (Fang-Yen, Avery, and Samuel 2009), or luciferin (Rodríguez-Palero et al. 2018) as a proxy. The ingestion of these artificial molecules can occur in the absence of bacterial food (Kiyama, Miyahara, and Ohshima 2012; Rodríguez-Palero et al. 2018), which may be seen as a disadvantage or an advantage depending on the application. For example, if the research question requires measuring intake without the complication of bacterial multiplication and metabolism, then a proxy may be preferred.

The reduction in bioluminescence that results from naloxone treatment illustrates both a limitation and a possible advantage of using a signal that requires active bacterial metabolism. On the one hand, if the signal is completely abolished then measurement is impossible. On the other hand, knowing that a given treatment affects bacterial physiology may be useful information in interpreting any observed feeding differences, since drug effects on bacteria are known to also affect host physiology (Cabreiro et al. 2013; Scott et al. 2017; García-González et al. 2017). Another limitation of our method is its sensitivity: we were unable to detect single worm feeding, although this is likely to be possible using a higher magnification imaging system.

Wild *C. elegans* strains aggregate and feed in groups when grown in the lab much like the *npr-1* mutants studied here, while the N2 laboratory reference strain are solitary feeders (de Bono and Bargmann 1998). The most commonly cited hypothesis to explain why wild isolates aggregate is that aggregation is useful to avoid high oxygen environments that represent oxidative stress, UV damage and desiccation risks (Busch and Olofsson 2012; Rogers et al. 2006). Based on our observations that DA609 worms clear bacteria inside clusters before moving onto new regions of the lawn (Figure 3C-D), and that in larger swarms only the leading edge is in contact with bacteria (Figure 4E-F), we hypothesise that collective feeding may in fact be a kind of hygienic behaviour in *C. elegans* that could reduce the risk of infections occurring through cuticle attachment.

## MATERIALS AND METHODS

### Reagent Table

| Resource | Designation | Source or reference | Identifiers | Additional Information |
| --- | --- | --- | --- | --- |
| Strain (*C. elegans*) | N2 | *Caenorhabditis* Genetics Centre | RRID:WB-STRAIN:N2 | Laboratory reference strain |
| Strain (*C. elegans*) | DA1116 | *Caenorhabditis* Genetics Centre | RRID:WB-STRAIN:DA1116 | Genotype: *eat-2(ad1116)ll*. |
| Strain (*C. elegans*) | DA609 | *Caenorhabditis* Genetics Centre | RRID:WB-STRAIN:DA609 | Genotype: *npr-1(ad609)X*. |
| Strain (*E. coli*) | DH5α-ilux | Addgene | RRID:Addgene_107879 | ilux pGEX(-) |
| Strain (*E. coli*) | OP50-GFP | Jonathan Hodgkin (University of Oxford) | RRID:WB-STRAIN:OP50-GFP |  |
| Strain (*E. coli*) | OP50-DsRed | Jonathan Hodgkin (University of Oxford) |  |  |
| Equipment | IVIS Spectrum In Vivo Imaging System | PerkinElmer | 124262 |  |
| Software, Algorithm | Living Image software | PerkinElmer | RRID:SCR_014247 | Version 4.3.1 |
| Chemical compound, drug | serotonin | Sigma-Aldrich | H7752 |  |
| Chemical compound, drug | naloxone | Sigma-Aldrich | PHR1802 |  |

### *C. elegans* maintenance and synchronisation

*C. elegans* strains used in this study are listed in the Reagents Table above. All worms were grown on *E. coli* OP50 at 20°C as mixed stage cultures under uncrowded and unstarved conditions, and maintained as described (Brenner 1974). Synchronised young adult animals were used for all imaging experiments, and they were obtained by bleach-synchronisation and subsequent re-feeding of starved L1’s on OP50 for 65-72 hours at 20°C.

### Measure 40 worm feeding with bioluminescent or fluorescent bacteria

A step-by-step protocol can be found at: <u>dx.doi.org/10.17504/protocols.io.5hsg36e</u>.

For every set of experiments, a fresh overnight liquid culture of DH5α-ilux or OP50-GFP was grown by inoculating a single bacterial colony into 100 mL of LB broth containing 50 μg/mL ampicillin and incubating overnight at 37°C at 220 rpm. The liquid culture was allowed to cool down to room temperature for 3-6 hours before use. 20 μL of the liquid culture was seeded onto the centre of a 35 mm low peptone (0.013% w/v) NGM plate and dried in a laminar flow hood (Heraguard) for 0.5 hour. Synchronised young adult worms were harvested and washed in M9 buffer, and 40 animals were transferred onto the seeded plate using a glass pipette without disturbing the bacterial lawn. After M9 was absorbed into the media, the imaging plate was gently vortexed for 10 seconds on the lowest setting of a vortex mixer (Vortex-Genie 2, Scientific Industries) to randomise initial worm positions. Imaging acquisition commenced 1 minute after the vortex start using the IVIS Spectrum imaging system (Caliper LifeSciences) and Living Image software (v 4.3.1). For bioluminescence, 1 second exposures were used with blocked excitation and open emission filters; for fluorescence, 0.5 second exposures were used with 465 nm excitation and 520 nm emission filters. Images were acquired every 6 minutes for up to 13.5 hours at 20°C and raw signals from user-defined regions of interest were extracted using Living Image software for downstream analysis. Field-of-view option C (13.5 cm × 13.5 cm) was used to allow simultaneous imaging of up to nine 35 mm plate feeding samples in 3×3 configuration, where at least one sample is a no-worm control to enable subsequent signal normalisation.

### Measure feeding during large population swarming with bioluminescent or fluorescent bacteria

A step-by-step protocol can be found at: <u>dx.doi.org/10.17504/protocols.io.53kg8kw</u>.

The bacteria overnight liquid culture was grown as described above. 500 μL of the liquid culture was seeded onto the centre of a 90 mm low peptone (0.013% w/v) NGM plate and dried in a laminar flow hood (Heraguard) for 2.5 hours. A separate 20 μL liquid culture was seeded onto the centre of a 35 mm low peptone plate and dried in a laminar flow hood (Heraguard) for 0.5 hours to serve as a no-worm control. Synchronised young adult worms were harvested and washed in M9 buffer, and a few thousand animals were transferred onto the seeded 90 mm plate using a glass pipette without disturbing the bacterial lawn. Imaging acquisition commenced immediately after worm transfer using the IVIS Spectrum imaging system (Caliper LifeSciences) and Living Image software (v 4.3.1). For bioluminescence, 1 second exposures were used with blocked excitation and open emission filters; for fluorescence, 1 second exposures were used with 465 nm excitation and 520 nm emission filters. Images were acquired every 2 minutes for up to 4.5 hours at 20°C, and raw signals from user-defined regions of interest were extracted using Living Image software for downstream analysis. Field-of-view option C (13.5 cm × 13.5 cm) was used to allow simultaneous imaging of one 90 mm plate swarming sample and one 35 mm plate no-worm control, the latter of which was used for subsequent signal normalisation.

### Measure 40 worm feeding after drug treatments

A step-by-step protocol can be found at: <u>dx.doi.org/10.17504/protocols.io.53ng8me</u>.

The protocol is essentially the same as the 40 worm feeding measurement protocol above, except for two differences: 1) Imaging plates are now low peptone NGM plates also containing drugs (20 mM serotonin or 10 mM naloxone), and 2) Young adult N2 worms were pre-starved on an unseeded NGM plate for 1 hour before being transferred onto seeded drug plates, and imaging commenced following a 1-hour drug exposure instead of immediately following worm transfer.

Drug plates were freshly prepared the day before each experiment. For serotonin (H7752, Sigma-Aldrich) treatment, low peptone NGM agar was prepared and serotonin was added to molten agar to a final concentration of 20 mM before the agar was dispensed into 35 mm plates. For naloxone (PHR1802, Sigma-Aldrich), a 10x stock solution was freshly prepared in water and 300 μL was spread on top of a 35 mm plate containing 3 mL low peptone NGM agar to achieve a final concentration of 10 mM. The naloxone plates were dried in a laminar flow hood (Heraguard) for 3 hours before all drug plates were wrapped in foil and stored at 4°C overnight for immediate use the next day.

To track worm positions following drug treatments, bright field imaging was performed using a custom-built six-camera rig equipped with Dalsa Genie cameras (G2-GM10-T2041) rather than the IVIS Spectrum imaging system. One-hour recordings were performed with 630 nm LED illumination (CCS Inc) at 25 Hz using Gecko software (v2.0.3.1), and worm positions were extracted from the pixel data using a MATLAB script.

### Measure pharyngeal pumping after drug treatments

Drug plates were prepared as described above, and were used either unseeded or seeded with 20 μL of DH5α-ilux overnight liquid culture. Pre-starved N2 young adult worms were transferred to drug plates with or without food as described above, and were exposed to the drugs for 1 hour before pharyngeal pumping was assessed. The number of pumps was scored over 60 seconds under a stereomicroscope (Zeiss Stemi 508).

### Data Analysis

Bioluminescence or fluorescence raw signals (photons/s) from imaging data were extracted from user-defined regions of interest using Living Image software (v 4.3.1). For each feeding experiment, the signal time series was divided by the level detected in the first frame. This relative signal was further normalised by the value of the corresponding no-worm control at each time point to correct for the non-stationarity of the signal in the absence of feeding. Relative feeding rates were then estimated by taking the derivative of the normalised signals over time.

## Data Availability

Supplementary Material is available on figshare. Strains and plasmids are available upon request. The authors affirm that all data necessary for confirming the conclusions of the article are present within the article and figures.

## Acknowledgements

ilux pGEX(-) was a gift from Stefan Hell (Addgene plasmid #107879; http://n2t.net/addgene:107879; RRID:Addgene_107879). *E. coli* OP50-DsRed and OP50-GFP strains are gifts from Jonathan Hodgkin. Some worm strains were provided by the CGC, which is funded by NIH Office of Research Infrastructure Programs (P40 OD010440). We thank Alex Sardini for assisting with the IVIS Spectrum imaging system.

## Author contributions

AEXB and KSS conceived the project. SSD designed and performed the experiments and conducted data analysis. The manuscript was written by SSD and revised by all authors.

