## Supplementary Material for "Measuring *C. elegans* spatial foraging and food intake using bioluminescent bacteria"

**Running title**: *C. elegans* feeding

**Key words**: bioluminescence, imaging, *C. elegans*, foraging, food distribution, feeding rate

**Corresponding Author**:

Andre E. X. Brown

Institute of Clinical Sciences
Imperial College London
Du Cane Road
London W12 0NN
United Kingdom

+44 (0)20 3313 8218


### Supplementary Movie Captions

Supplementary Movies can be viewed at: https://tinyurl.com/y2rml8vm.

**Supplementary Movie S1:** 40-worm feeding experiments with DH5α-ilux bioluminescent bacteria. Top three samples are DA609 worms, middle three samples are N2 worms, bottom left and middle samples are DA1116 worms, bottom right sample is a no-worm control. The movie plays at 10800x real speed.
**Supplementary Movie S2:** Large population N2 swarming experiment with DH5α-ilux bioluminescent bacteria. The top right sample is a no-worm control. The movie plays at 1800x real speed.
**Supplementary Movie S3:** Large population N2 swarming experiment with OP50-GFP fluorescent bacteria. The top right sample is a no-worm control. The movie plays at 3600x real speed.
**Supplementary Movie S4:** Large population DA609 swarming experiment with DH5α-ilux bioluminescent bacteria. The top right sample is a no-worm control. The The movie plays at 3600x real speed.

### Supplementary Figures

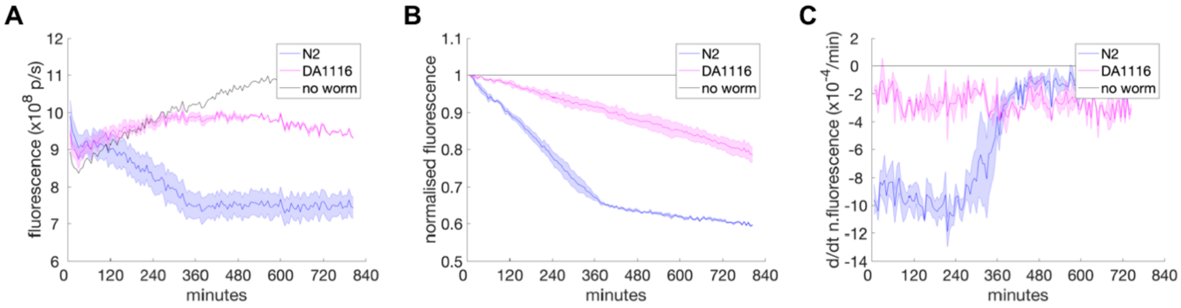


**Supplementary Figure S1**. Fluorescence signal from population feeding experiments of N2 and DA1116 worms, showing **A**) raw signal, **B**) normalised signal (normalised against the starting signal and then against the no-worm control signal), **C**) derivative of the normalised signal calculated over a 60-minute window. Forty N2 (blue) worms, forty DA1116 (magenta) worms, or no-worm control (black) experiments were performed on a 20 µL OP50-GFP lawn. 0.5 second exposure measurements were read every 6 minutes using 465 nm excitation and 520 nm emission filters. All samples shown were imaged simultaneously. Here n = 3 for N2, n = 2 for DA1116, n = 1 for control; error bars represent ±1 SD.

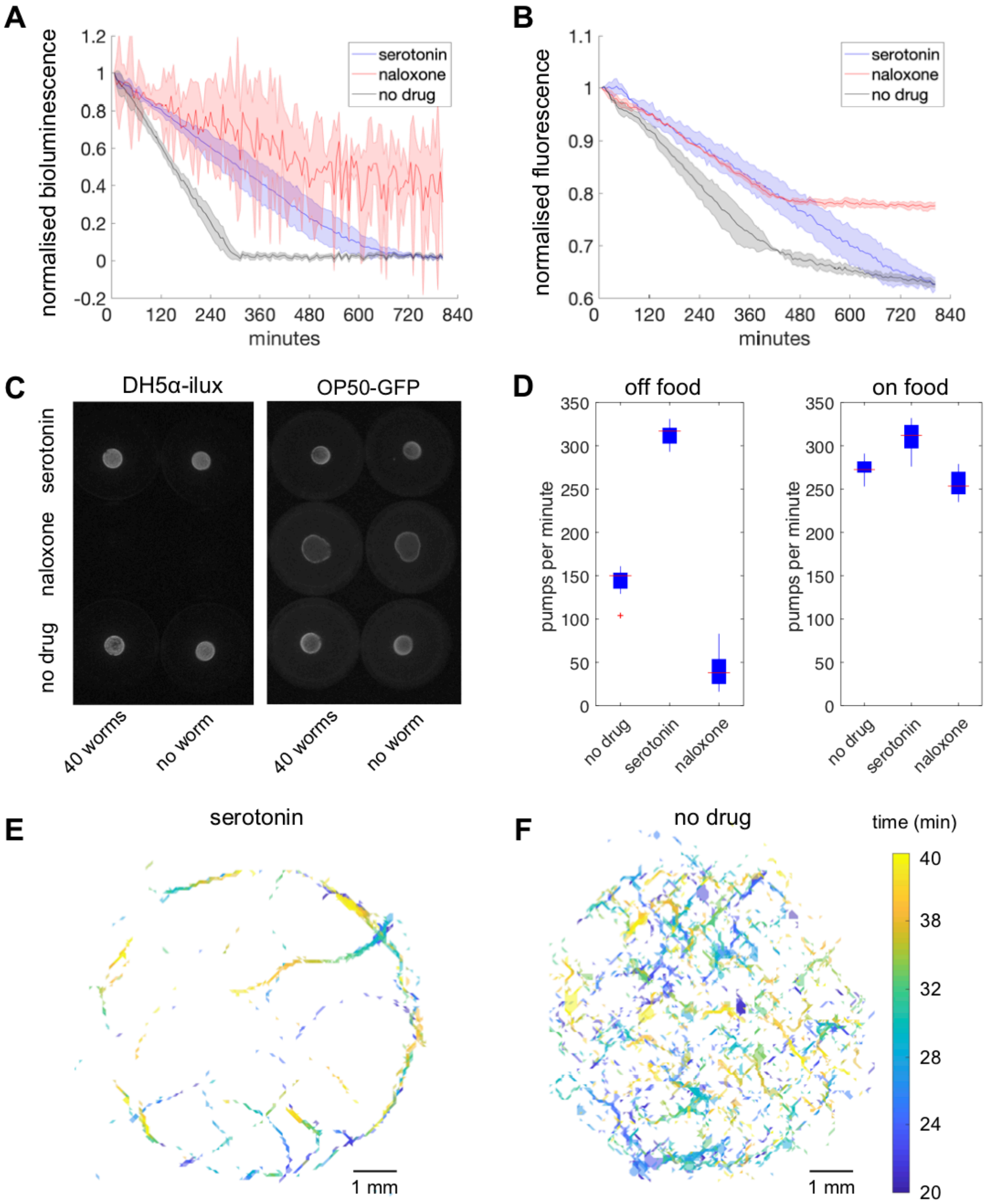


**Supplementary Figure S2**. The effect of drug treatments on *C. elegans* feeding rates and foraging. Forty pre-starved N2 worms were subject to serotonin-, naloxone-, or no-drug mock treatment for 1 hour on food before feeding was measured. **A-B**) Normalised signal from experiments on DH5α-ilux (A) or OP50-GFP (B) bacterial lawn following treatment with serotonin (blue), naloxone (red), or no-drug mock control (black). For A), 1 second exposure measurements were read every 6 minutes from the start of re-feeding. n = 4 for each condition pooled between two independent sets of experiments, error bars represent ±1 SD. For B), 0.5 second exposure measurements were read every 6 minutes from the start of re-feeding using 465 nm excitation and 520 nm emission filters. n = 2 for each condition, error bars represent ±1 SD. **C**) Snapshots of the signal level at the end of 1-hour drug exposure, which is the start of the feeding measurements. **D**) Pharyngeal pumps per minute of mock-, serotonin-, and naloxone-treated worms. Drug treatment was performed either off (left) or on (right) food, and pumping was scored for 60 seconds. n = 9 for no-drug off food, n = 11 for serotonin off food, n = 5 for naloxone off food, n = 10 for no-drug on food, n = 10 for serotonin on food, n = 10 for naloxone on food. **E-F**) Worm positions on the circular bacterial lawn during the 20-40 min imaging window following serotonin- (E) or mock- (F) treatment. Different colours indicate time progression according to the colour bar.

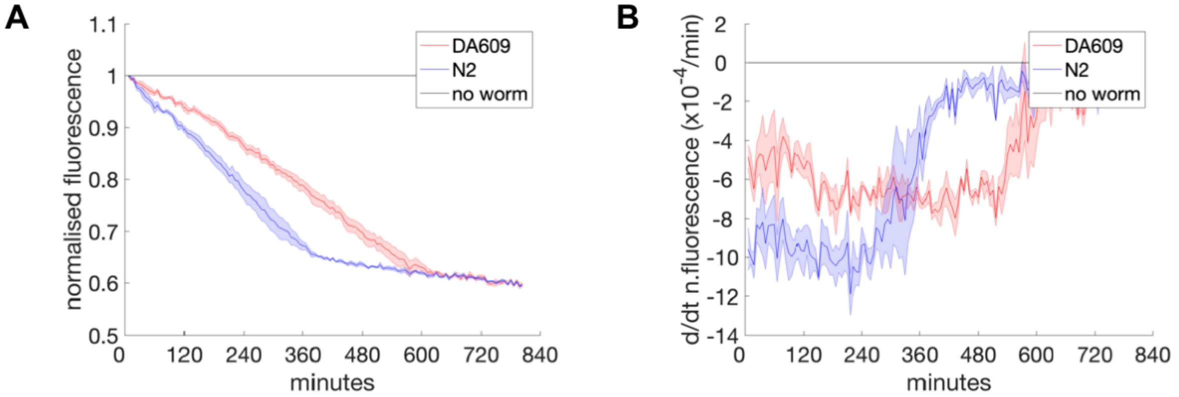


**Supplementary Figure S3**. Fluorescence signal from population feeding experiments of N2 and DA609 worms, showing **A**) normalised signal, and **B**) derivative of the normalised signal calculated over a 60-minute window. Forty DA609 (red) or N2 (blue) worms or no-worm control (black) experiments were performed on a 20 µL OP50-GFP lawn. 0.5 second exposure measurements were read every 6 minutes using 465 nm excitation and 520 nm emission filters. All samples were imaged simultaneously.  n = 3 for each condition; error bars represent ±1 SD.

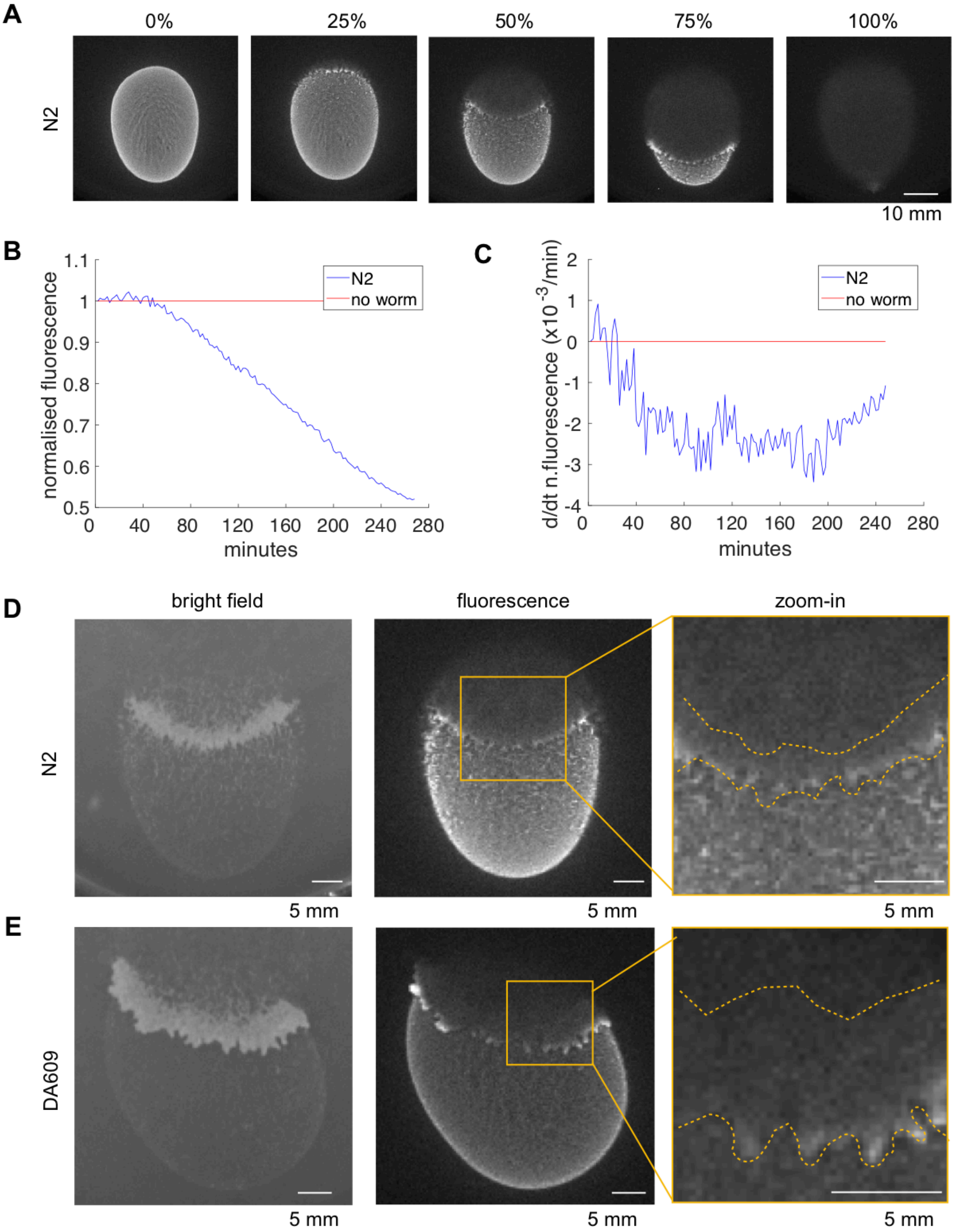


**Supplementary Figure S4**. Fluorescence signal from large population swarming experiments. A few thousand age-synchronised worms were allowed to feed and swarm over a 500 µL OP50-GFP lawn. **A**) Snapshots of N2 swarming experiments, with time progression to total food depletion indicated at the top. **B)** Normalised signal and **C)** derivative of the normalised signal calculated over a 20-minute window, from an N2 swarming experiment with OP50-GFP lawn. **D-E**) Sample snapshots from N2 (D) and DA609 (E) swarming experiments, showing bright field (left) and fluorescence (middle) channels. The boxes in the middle panels are zoomed in and displayed on the right, with the worm front outline shown in dashed yellow lines.
